# PI(4,5)P_2_ Binding Sites in the Ebola Virus Matrix Protein Modulate Assembly and Budding

**DOI:** 10.1101/341248

**Authors:** Kristen A. Johnson, Melissa R. Budicini, Sarah Urata, Nisha Bhattarai, Bernard S. Gerstman, Prem P. Chapagain, Sheng Li, Robert V. Stahelin

## Abstract

Ebola virus (EBOV) causes sever hemorrhagic fever in humans, can cause death in a large percentage of those infected, and still lacks FDA approved treatment options. In this study, we investigated how the essential EBOV protein, VP40, forms stable oligomers to mediate budding and assembly from the host cell plasma membrane. An array of *in vitro* and cellular assays identified and characterized two lysine rich regions that bind to PI(4,5)P_2_ and serve distinct functions through the lipid binding and assembly of the viral matrix layer. We found that when VP40 binds PI(4,5)P_2_, VP40 oligomers become extremely stable and long lived. Together, this work characterizes the molecular basis of PI(4,5)P_2_ binding by VP40, which stabilizes formation of VP40 oligomers necessary for viral assembly and budding. Quercetin, a natural product that lowers PI(4,5)P_2_ in the plasma membrane, inhibited budding of VP40 VLPs and may inform future treatment strategies against EBOV.

## Introduction

Ebola virus (EBOV) is a lipid-enveloped pleomorphic virus that can infect a wide variety of human cells causing significant disease^1,2^. To date there are no FDA approved drugs or vaccines for EBOV treatment and there is still a great deal of need in understanding how to therapeutically target the five different types of *ebolaviruses*. The EBOV glycoprotein (GP), which studs the exterior of the virus particle during the infection scheme, has been the major target of vaccine development and antibody based therapies. Indeed, a vaccine using EBOV GP pseudotyped into vesicular stomatitis virus is being implemented in the current EBOV outbreak in the Democratic Republic of Congo^3^. While many of the emerging treatments aimed at the GP show great promise, the GP has been shown to mutate significantly during the course of the recent outbreak^4-6^ and a watch list of potential GP antibody escape mutants has been generated^7^. Thus, other targets in the viral life cycle are being explored such as host-viral interactions^8^ and the matrix protein VP40^8-10^.

VP40 is the matrix protein of EBOV and required for viral assembly, budding and viral spread^11-16^. As the most abundantly expressed EBOV protein, VP40 achieves several distinct and essential tasks to ensure viral success. VP40 is a transformer protein with several structures known, including a monomer^17^, dimer, hexamer and octamer^18^. Initially, VP40 enters the nucleus of the host cell post viral entry^19^, likely binds RNA as an octamer and regulates viral genome replication^18,20.^ Subsequently, VP40 dimers interact with host lipids such as phosphatidylserine (PS)^21,22^ to localize to the plasma membrane (PM)^18,22^ and form stable hexamers^18,21,23-25^. Phosphatidylinositol 4,5-bisphosphate (PI(4,5)P_2_) at the host PM inner leaflet is required for large VP40 oligomers to form^26^ from PS induced VP40 hexamers^18,21-25^.

Despite an important role for PS and PI(4,5)P_2_ in the EBOV life cycle and budding from the host cell PM, the molecular basis and structural consequences of VP40-lipid interactions are mostly unknown. Notably, the VP40 dimer is necessary for PM localization whereas the monomer is not sufficient for trafficking to the PM^18,22^. Dimer interactions with PM lipids are purported to induce structural changes to form VP40 hexamers^18,21,23-25,27^ and larger oligomers forming the viral matrix layer^23,26^. Notably, the VP40 octamer has been shown to play an important role in the regulation of viral transcription^18^ and has not been detected at the PM nor in virions or VLPs. Additionally, the VP40 octamer was found to have a significant reduction in PS affinity compared to the VP40 dimer^22^.

Previously, we found that VP40 requires PI(4,5)P_2_ at the plasma membrane for the formation of large self-oligomers, VLP formation, and VLP budding^26^. A recent molecular dynamics model found VP40 hexamers cluster PI(4,5)P_2_ but not PS at the PM through C-terminal lysine residues^28^. In this study, we investigate the VP40 lipid binding properties at the PM and the structural consequences of interactions with PS and PI(4,5)P_2_. Results demonstrate that PI(4,5)P_2_ provides significant structural stability to VP40 hexamers that form following interactions with PS. PI(4,5)P_2_ provides extensive stability to the viral matrix layer through two distinct PI(4,5)P_2_ binding sites, loss of which leads to rapid dissociation of VP40 oligomers and inhibition of viral budding. Further, a natural product Quercetin, which reduces plasma membrane PI(4,5)P_2_ significantly inhibited VLP formation from the PM. Taken together, these new findings account for the molecular basis of PI(4,5)P_2_ recognition by VP40, PI(4,5)P_2_ induced structural stability and assembly of VP40, and open up a potentially new avenue for EBOV treatment.

## Results

### Hypothesized PI(4,5)P_2_ binding site is conserved among *ebolaviruses*

VP40 from Zaire, Sudan, Tai Forest, Bundibugyo, and Reston *ebolaviruses* were aligned using uniprot to reveal sequence conservation in residues hypothesized to bind to PI(4,5)P_2_. Lys^224^, Lys^225,^ Lys^274^, and Lys^279^ are completely conserved while Lys^221^ has an Arg in the Tai Forest strain (Figure 1A). Lys^225^ is Arg in the Bundibugyo strain while residue 270 is Arg or Lys (Figure 1A). Both Arg and Lys are commonly found in PI(4,5)P_2_ binding sites^29^. To test if these VP40 residues are involved in PI(4,5)P_2_ binding, alanine scanning mutagenesis was used for C-terminal domain regions with high conservation across *eboaviruses*. These mutations include Lys^104^, Lys^221^, Lys^224^, Lys^225^, Lys^236^, His^269^, Lys^270^, Lys^274^, Lys^275^, Lys^279^, Lys^291^. When mapped onto the VP40 hexamer model, the 220’s lysine residues (blue) and the 270’s lysine residues (green) formed distinct, adjacent domains (Figure 1B).

**Figure 1:**
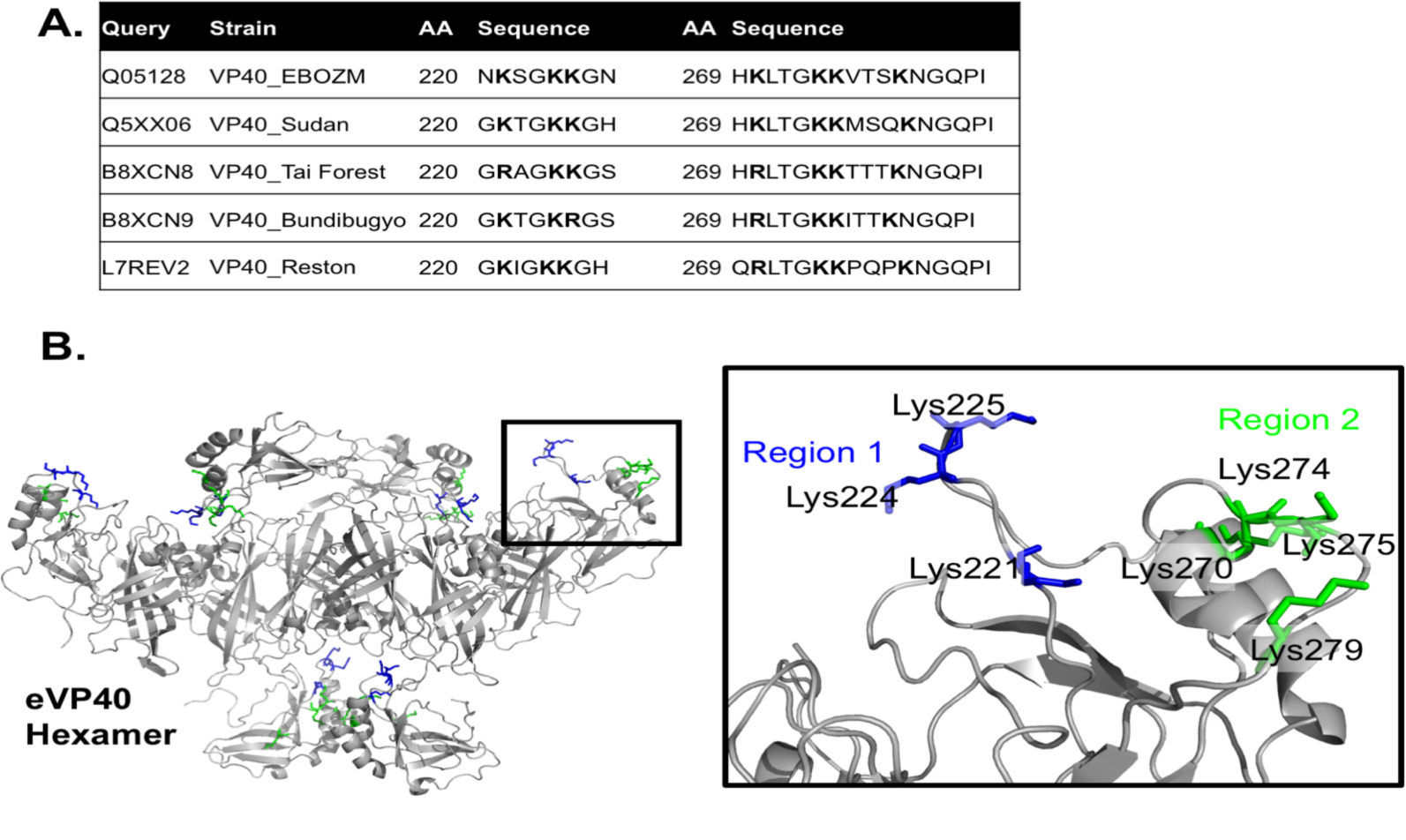
Ebola VP40 sequence and structure revels two PI(4,5)P_2_ binding sites. **A.** Consensus sequence of Ebola VP40 strains several lysine residues in the 220 and 270 regions of the protein are positioned at the surface of the C-terminal domain. Sequence alignment was performed with uniprot alignment tools. **B.** Zaire VP40 hexamer structure (4ldd with modeled C-terminal domains) with Region 1 (220 lysine residues) colored in blue and Region 2 (270 lysine residues) colored in green.

### VP40 binds to PI(4,5)P_2_ containing membranes with high selectivity among phosphoinositides

VP40 binding to phosphoinositides (PI(3)P, PI(4)P, PI(5)P, PI(3,4)P_2_, PI(3,5)P_2_, PI(4,5)P_2_, and PI(3,4,5)P_3_) was quantified using a raffinose pentahydrate loaded large unilamellar liposome (LUV) pelleting assay^30^. The liposome composition included 1,2-dipalmitoyl-*sn*-glycero-3-phosphocholine (DPPC) and cholesterol 1:1 with 5% phosphoinositide and 2% dansylPE. Previously, we did not observe binding to PI(4,5)P_2_ containing multilamellar vesicles (MLVs) without 40% POPS added to the PIP^31^. However, we found that VP40 binds well to PI(4,5)P_2_ in the LUV system, which better recapitulates cellular membrane bilayers. The binding to PI(4,5)P_2_ was especially notable in the presence of ordered lipids DPPC and cholesterol when compared to POPC and DOPE (See **Figure S1A and S1B**). To ensure LUV formation, dynamic light scattering was used to measure the diameter of extruded liposomes (data not shown). Pelleting efficiency was quantified by including 2% dansylPE in the liposomes, the percent fluorescence in the supernatant (SN) and pellet (P) fractions were quantified (Ex 355, Em 510). Pelleting efficiency for this liposome composition was approximately 100% (**Figure S1C**).

In this LUV system, VP40 displayed significant binding to PI(4)P, PI(3,4)P_2_, and PI(4,5)P_2_ with the most significant binding to PI(4,5)P_2_. Interestingly, binding to PI(3,5)P_2_ was not observed (Figure 2A and 2B). VP40 selectivity for PI(4,5)P_2_ over the other 6 phosphoinositide species indicated there was likely a lipid binding site driving the selectivity and additionally, previous studies ruled out nonspecific electrostatics as a driving force of EBOV VP40 assembly^21,26^.

**Figure 2:**
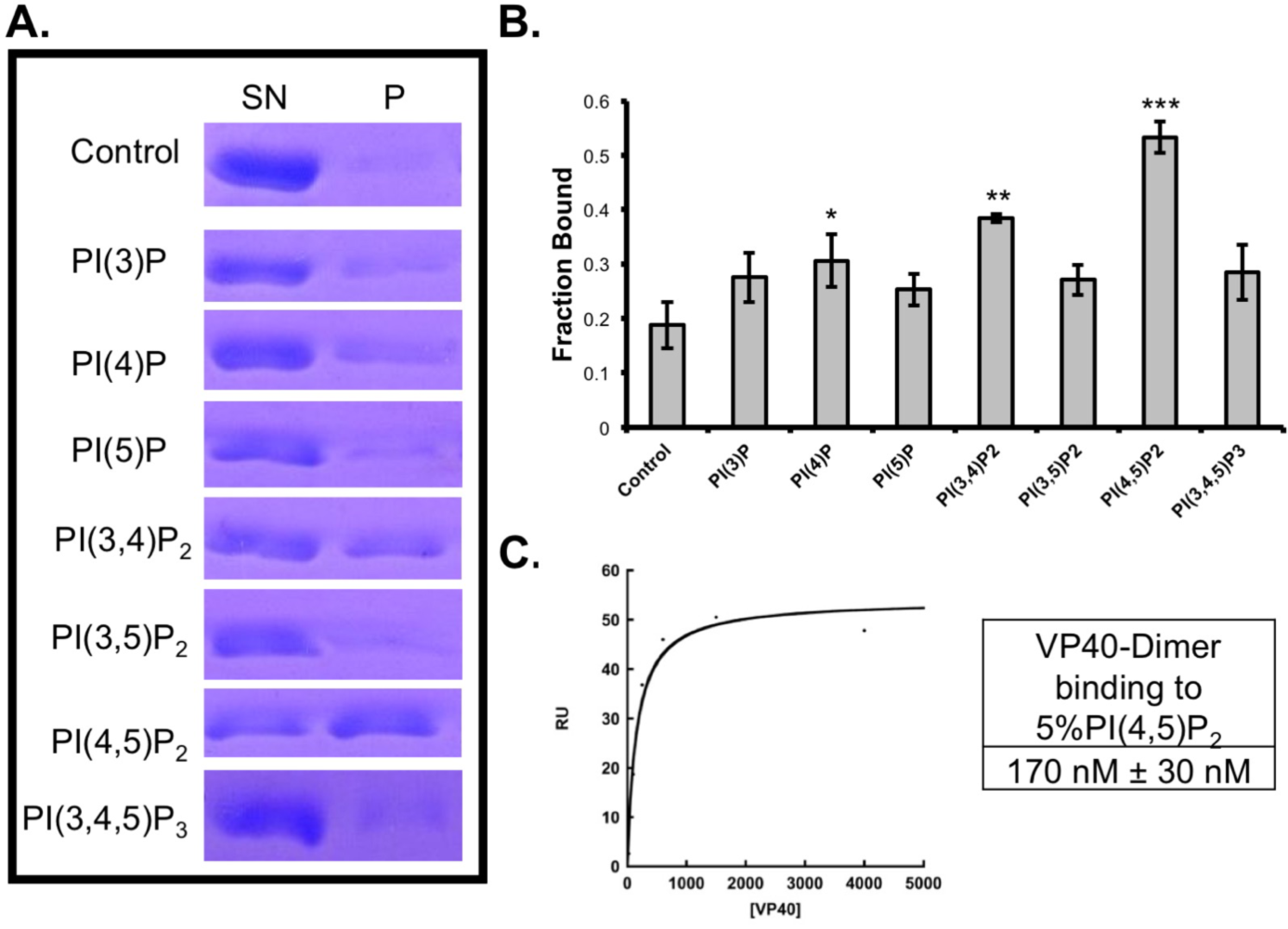
VP40 binds to PI(4,5)P2 with selectivity and nanomolar affinity. **A.** Representative SDS PAGE bands from the large unilamellar (LUV) pelleting assay with DPPC:Cholesterol without (control) or with 5% of one of the seven phosphatidylinositol phosphate species. **B.** Quantified VP40 fraction bound to liposomes (protein found in pellet fraction). The average value is plotted and shown ± the SEM. **C.** Representative surface plasmon resonance curve of VP40 binding to liposomes containing 5% PI(4,5)P_2_ gave an average binding affinity of 170 nM with a standard error of the mean of 26 nM. * P<0.05, ** P<0.001, ***P<0.00001

Surface plasmon resonance (SPR) was used to determine the affinity of VP40 for PI(4,5)P_2_ as previously described^32^. Using this technique, we found that the VP40 dimer binds to PI(4,5)P_2_ containing LUVs with an apparent affinity of 170 nM (Figure 2). This binding affinity is also slightly stronger than the interaction between VP40 and POPS, also determined with SPR^21^.

### VP40 binds to PI(4,5)P_2_ through lysine residues in CTD Region 1 and Region 2

To determine which VP40 residues are responsible for PI(4,5)P_2_ interactions, lipid binding of VP40 lysine mutants was tested *in vitro* using a the LUV assay detailed above. Binding to liposomes containing 5% PI(4,5)P_2_ was measured for the dimer of VP40-WT and select lysine mutants. VP40-K104A and K236A did not have a significant difference in binding to PI(4,5)P_2_ LUVs compared to the WT protein (Figure 3). In contrast, K221A, K224A, K225A, K270A, K274A, K275A, and K279A had significant decreases in binding to LUVs containing PI(4,5)P_2_ compared to WT (Figure 3).

**Figure 3:**
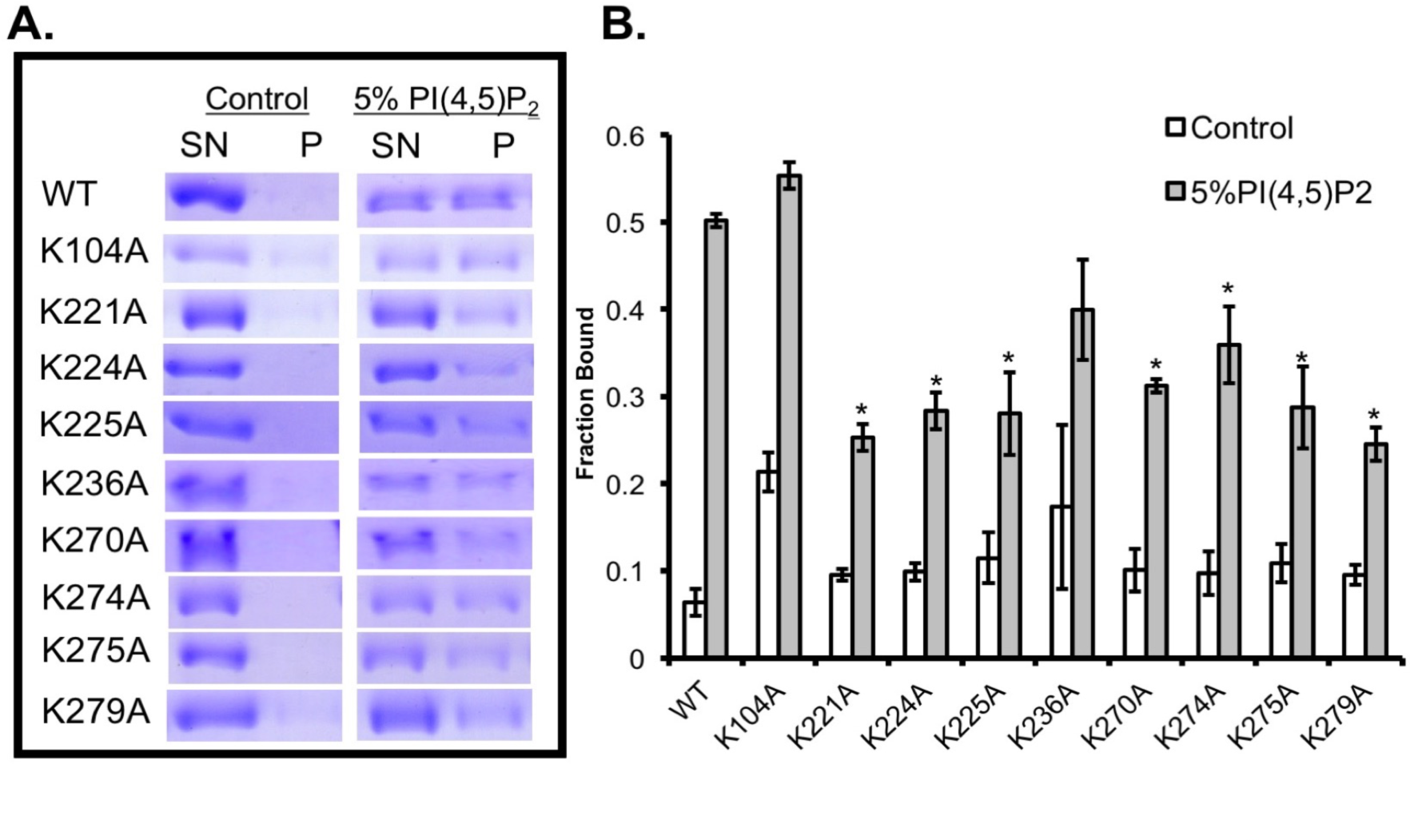
VP40 C-terminal domain lysine residues are important for PI(4,5)P_2_ binding. **A.** Representative SDS PAGE of the LUV pelleting assay to measure VP40 binding to control (DCCP and Cholesterol) or 5% PI(4,5)P_2_. **B.** Average fraction bound to LUVs shown ± the SEM, * P<0.05.

### PI(4,5)P_2_ binding residues are critical for the VP40 budding phenotype in live cells

In COS-7 cells, WT VP40-EGFP produces an abundance of VLPs at the PM 12-14 hours post transfection (Figure 4A)^26^. To screen the phenotype of VP40 C-terminal mutants, each EGFP-construct was expressed in COS-7 cells. The percentage of cells producing VLPs were compared to WT-VP40 (Figure 4B), representative images of each are shown in Figure 4A. Mutations that showed two phenotypes have a second, smaller representative image to showthe minor phenotype. K221A and K225A had the most dramatic reduction in VLP production with fewer than 10% of cells producing VLPs. Less than 20% of cells expressing K221F and K224A showed VLPs and about 50% of cells expressing K270A, K274A, or K275A had detectable VLPs. Some lysine to alanine mutations (K104A, K236A, K279A, K291A) had a phenotype similar to WT.

**Figure 4:**
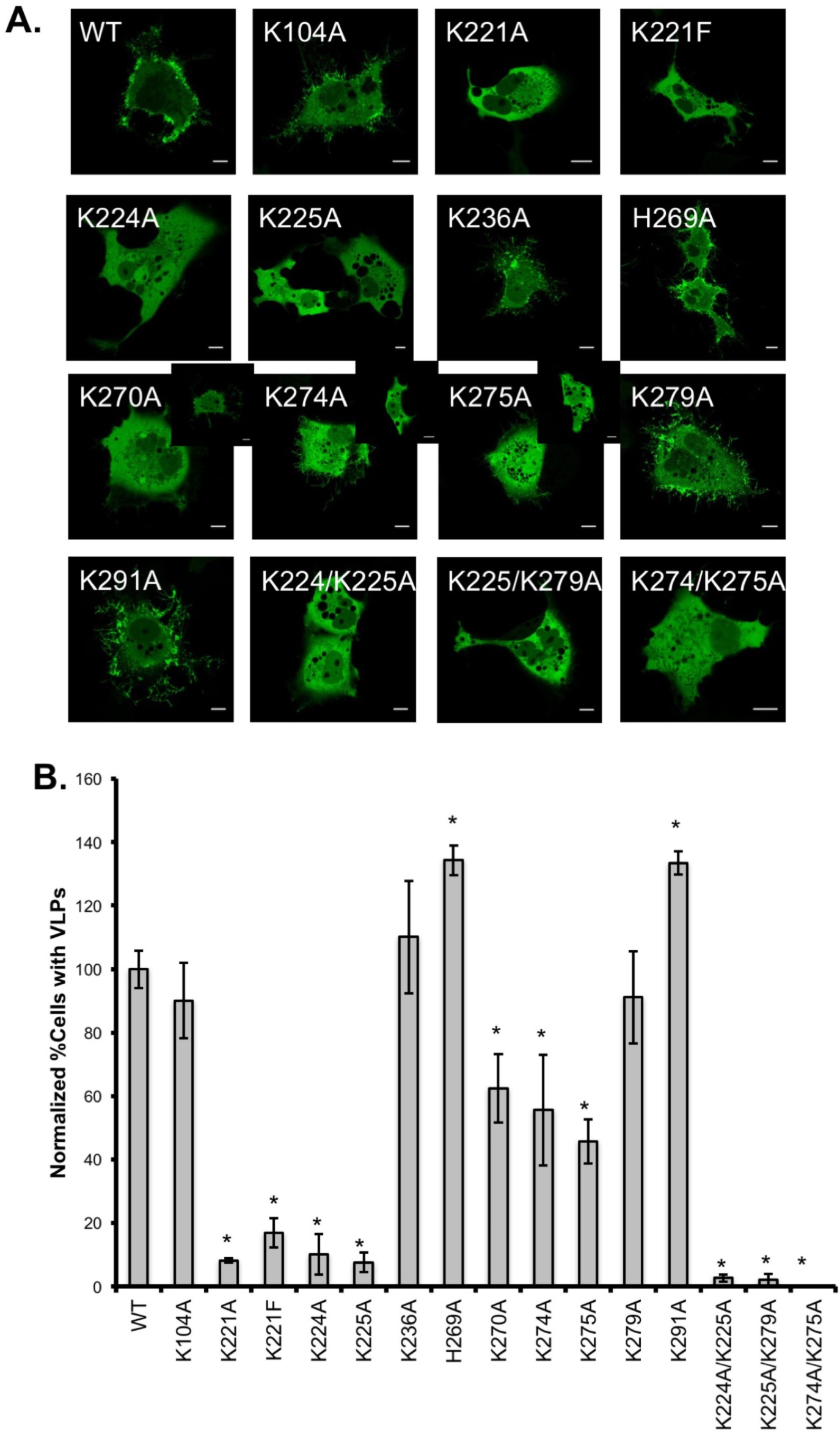
VP40 mutant phenotype in live cells. **A.** Representative images of WTVP40-EGFP and VP40 mutants 14 hours post transfection in COS-7 cells. Scale bars are 10 μm. **B.** Percentage of the cell population expressing VLPs 14 hours post transfection, normalized to WT. Average values are shown ± the SEM, * P<0.05.

Scanning electron microscopy (SEM) was used to achieve high resolution of the PM of VP40 transfected cells. VP40-EGFP-WT, K221A, K104A, K224A, K274A, K279A, and non-transfected COS-7 cells were prepared over two independent experiments (Figure 5B and 5C). COS-7 cells have naturally occurring filopodia structures observed in SEM (see **Figure S2**). Because of this, filopodia structures were measured in FIJI to determine the average control and VP40 transfected cell filopodia length. We found that control cells have an average of 1µm filopodia while VP40 WT transfected cells have closer to 2 µm VLPs (Figure 5). The VLP length was also compared to EGFP-VP40 from confocal imaging (Figure 5). In consonance with the confocal imaging, K104A and K279A had similar VLP structures to WT-VP40. K274A had similar length but less VLP density and K221A and K224A had filopodia structures more similar to the non-transfected control than WT-VP40 (See Figure 5).

**Figure 5:**
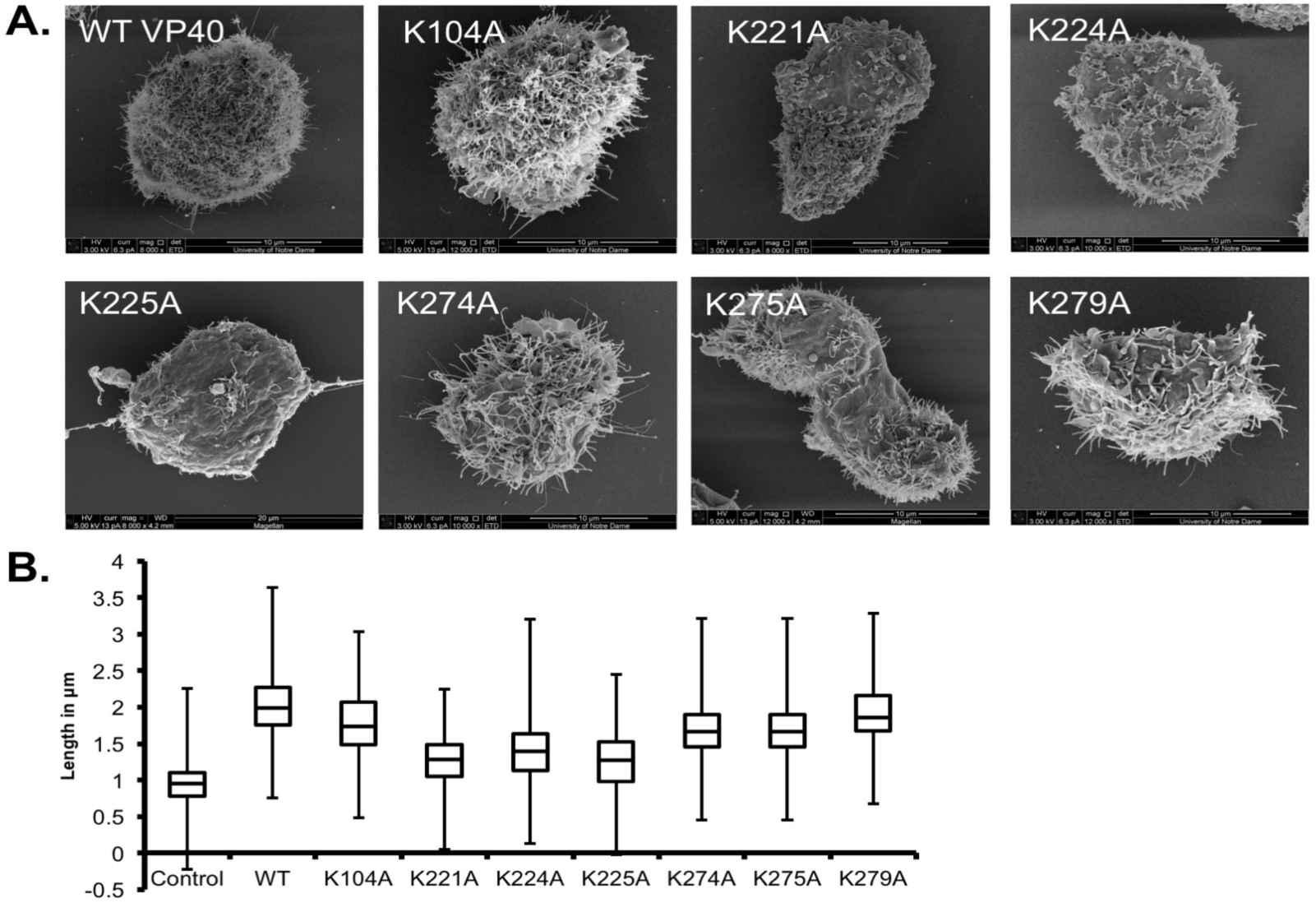
Scanning electron microscopy of VP40 expressing cells to assess virus like particle production. **A.** Representative SEM images of COS-7 cells expressing WT-VP40-EGFP and mutant-VP40-EGFP constructs. **B.** Box and whisker plot shows VLP lengths for WT and mutant VP40s. Values represent the average VLP length where 20 VLPs measured per image per mutant, data represents two independent experiments.

### PI(4,5)P_2_ binding resides in Region 1 are critical for VP40 oligomers larger than a hexamer

A VP40 oligomerization assay in live cells^21,23-26,33^ was used to compare oligomerization abilities of mutants deficient in PI(4,5)P_2_ binding compared to WT. Previously, when PI(4,5)P_2_ was depleted from the PM of live cells with a Myc5-PtaseIV construct, VP40 large oligomer formation was significantly reduced^31^. Adu-Gyamfi et al. found VP40 hexamer formation was dependent on plasma membrane PS^21^. Together, this suggests different roles for PS and PI(4,5)P_2_ in the structure and assembly of the viral matrix layer.

We assessed the oligomer formation of VP40 mutants and found that K221A, K224A, and K225A had a dramatic and significant reduction in oligomers larger than a hexamer (Figure 6). For these mutants, there was also a significant increase in the hexamer population compared to WT VP40. K270A, K274A, and K275A also had a significant decrease in oligomers larger than a hexamer but some large oligomers were detected (Figure 6). This data recapitulates what was observed previously when PI(4,5)P_2_ was constitutively depleted with Myc5’Ptase IV constructs^31^ but suggests the two cationic regions may contribute differently to the oligomerization and stabilization process.

**Figure 6:**
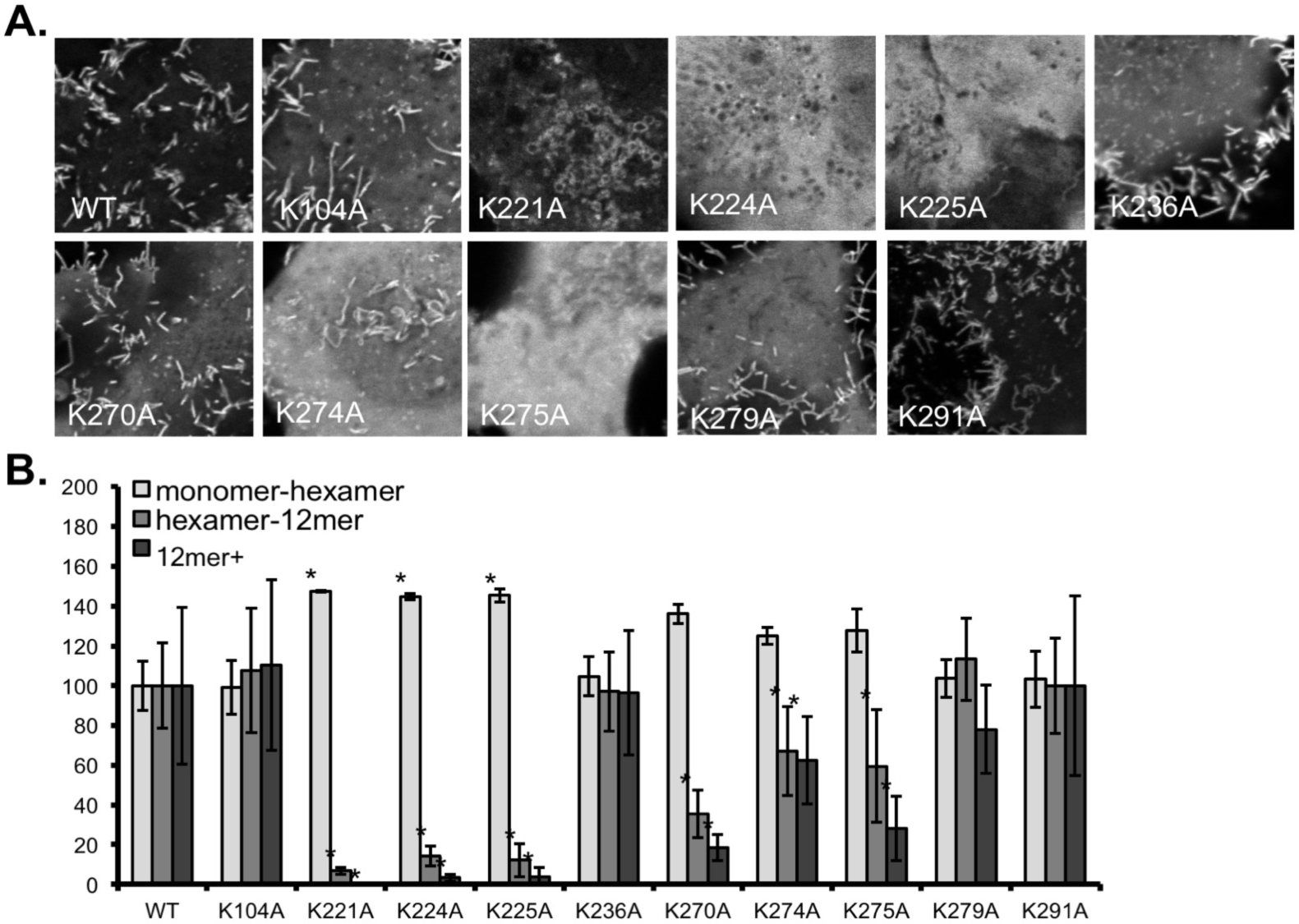
Number and Brightness analysis to determine VP40 oligomerization in live cells. **A.** Representative image frame from the 100 images acquired per for the imaging experiment. All are imaged at the same zoom (16.4). **B.** Normalized VP40 oligomer formation. The average WT-VP40-EGFP population was normalized to 100% for monomer-hexamer, hexamer-12mer, and 12mer+. WT and mutant VP40-EGFP oligomer population values are shown ± the SEM, * P<0.05.

### PI(4,5)P_2_ binding residues are critical for efficient virus like particle budding

Next, VLPs and cell lysates of VP40 transfected COS-7 cells were collected 20-24 hours post transfection (see Methods for more details). The budding efficiency of WT VP40 and select mutants was determined with western blot analysis. Surprisingly, K221A has a similar budding efficiency to WT while K221A expression levels were consistently lower than WT. K221A VLPs were not commonly observed in confocal microscopy (Figure 5B) or scanning electron microscopy (Figure 6B) and oligomer formation was significantly reduced (Figure 7B).

**Figure 7:**
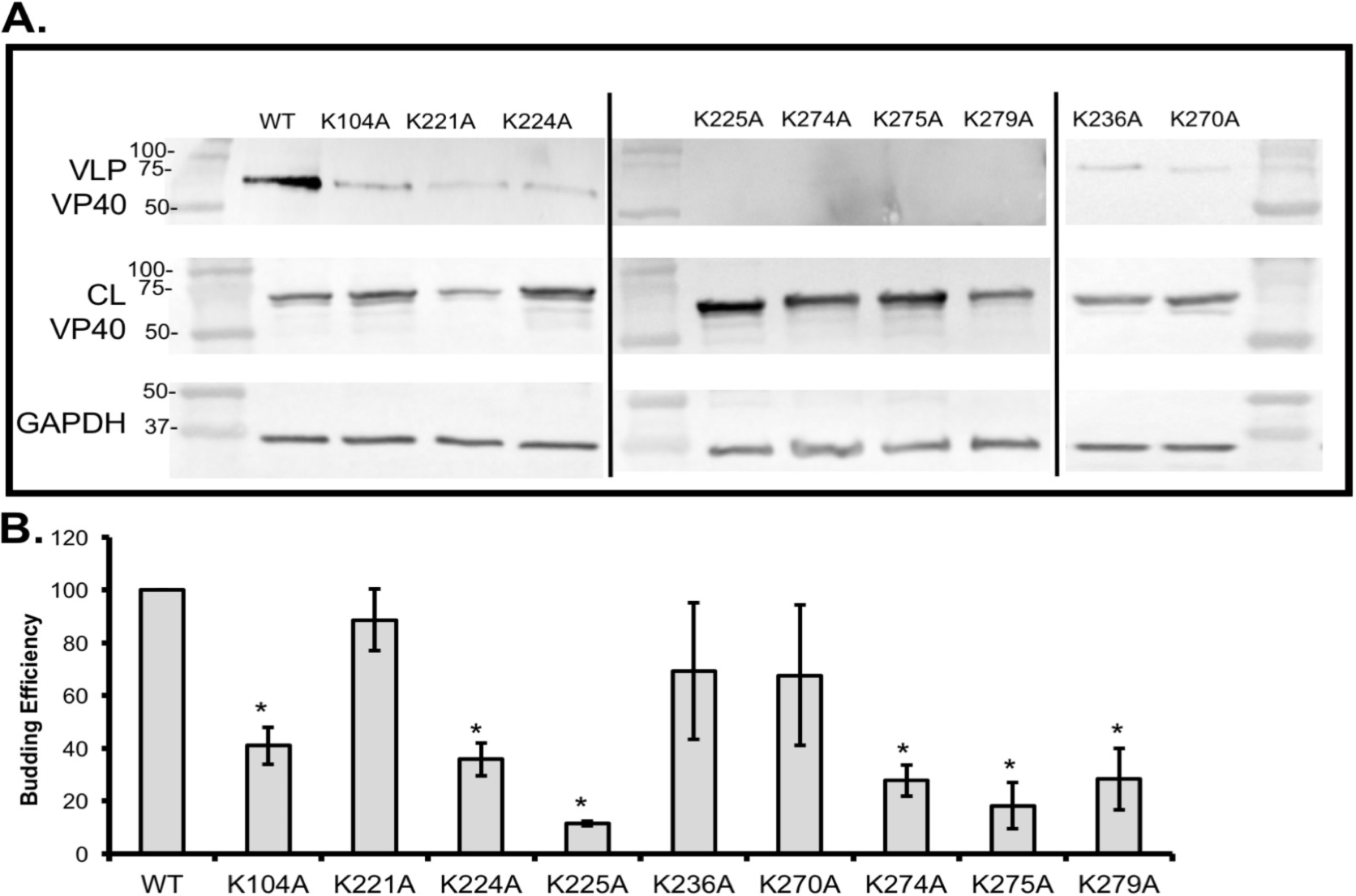
VLP budding efficiency of WT and mutant VP40. **A.** Western blots of WT and mutant VP40-EGFP show VP40 in the VLP and the cell lysate (CL) fractions. Housekeeping protein, GAPDH (CL), was used as a loading control for these experiments. Molecular weight markers are shown for all blots to verify band size for VP40-EGFP and GAPDH. **B.** Quantified budding efficiency normalized to VP40-WT. Average values are shown ± the SEM, * P<0.05.

K104A exhibited a decrease in VLP formation. This mutant is interesting as it is in the N-terminal domain but folds against the C-terminal domain adjacent to aspartic acid residue 193 in the dimer. This suggests its interaction with lipid may not be a major contributor to PI(4,5)P_2_ dependent assembly but a role in interdomain interactions cannot be discounted at this time. K279A also showed a significant decrease in budding which is consistent with the reduction in binding to PI(4,5)P_2_ containing membranes *in vitro* (Figure 3). This result was partially unexpected as VLPs are observed in confocal microscopy (Figure 4B) and SEM (Figure 5B). However, this mutation had a significant decrease in PI(4,5)P_2_ binding in the LUV pelleting assay (Figure 3B) and a small decrease in oligomerization (Figure 6). Because Lys^279^ is important for lipid binding *in vitro*, we hypothesize this indicates that even a small decrease in PI(4,5)P_2_ binding and self-oligomerization may have dramatic effects on budding efficiency.

### Hydrogen deuterium exchange mass spectrometry reveals VP40 oligomers become extremely stable following PI(4,5)P_2_ binding

Hydrogen deuterium exchange mass spectrometry (HDX MS) was used to better understand structural changes in VP40 upon binding to PS and PI(4,5)P_2_ containing liposomes. Complete coverage analysis of the VP40 protein was achieved with many overlapping peptides (**Figure S3**). To ensure a homogeneous sample, elevated levels of PI(4,5)P_2_ or PS were used (**Figure S4**). Samples were incubated for 10,100, 1000, 10,000 and 100,000 seconds (s). Changes in deuterium incorporation during these time points are shown in **Figures S5-10**. Supplemental figures 4, 6, and 8 show ribbon maps with the level of deuterium incorporation mapped to the protein sequence. Supplemental figures 5, 7, and 9 show the influence of PC, PS, and PI(4,5)P_2_, respectively. **Figure S10** shows the HDX data mapped onto the dimer structure and **Figure S11** shows the influence of each lipid on the VP40 dimer structure at 10,000 s. The deuterium incorporations for each time point and lipid were also mapped to the VP40 hexamer structure, the relevant structure for VP40 matrix assembly^18^.

**Figure 8.**
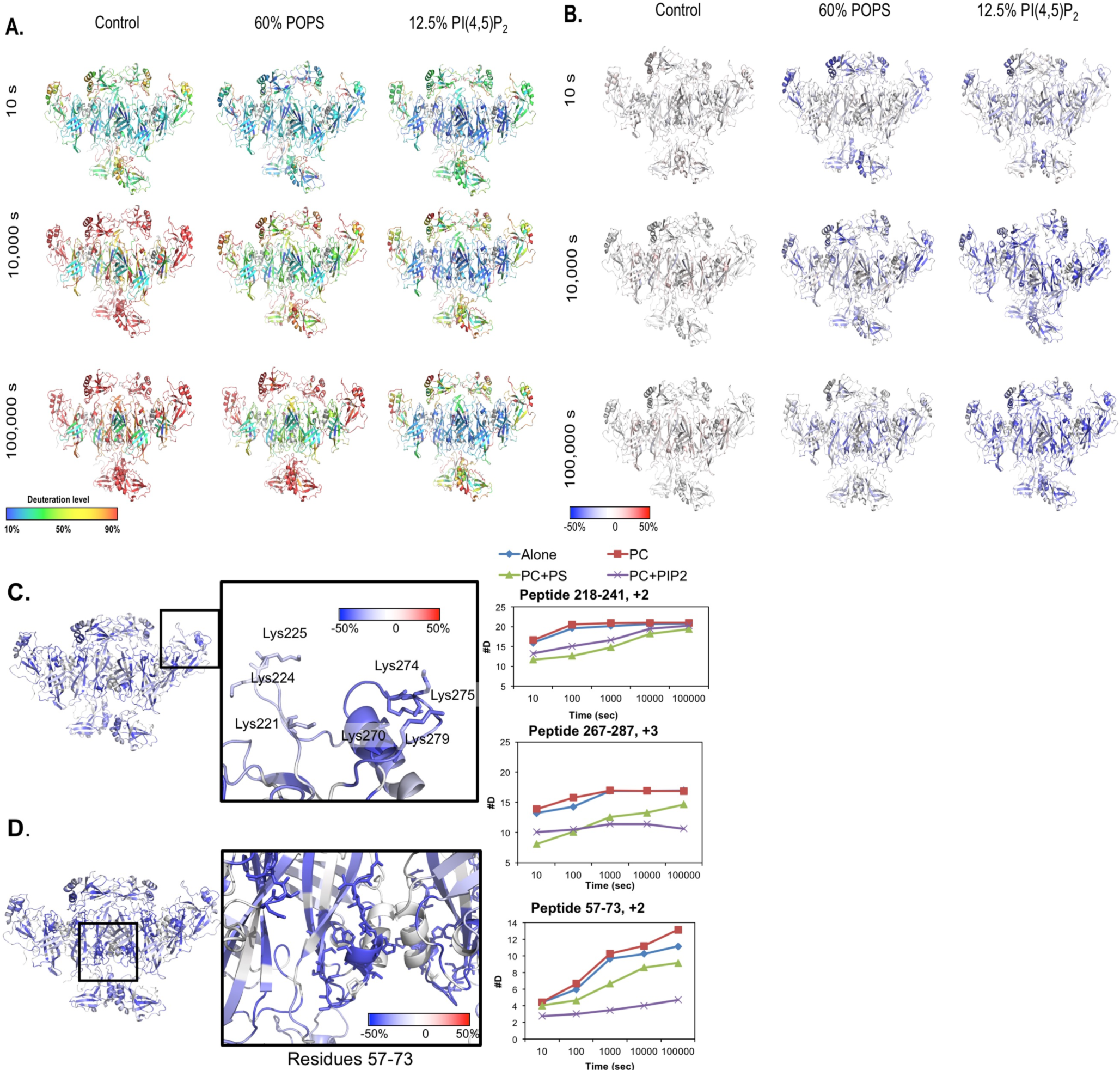
HDX-MS reveals changes in VP40 solvent accessibility when co incubated with various liposomes. **A.** Deuteration level of VP40 (shown as the hexamer) at 10 s, 10,000 s and 100,000 s time points after a 30 minute incubation with Control liposomes, PS containing liposomes, or PI(4,5)P_2_ containing liposomes. **B.** The influence of Control liposomes, PS containing liposomes, or PI(4,5)P_2_ containing liposomes on VP40 deuteration is mapped onto the VP40 hexamer structure for the 10 s, 10,000 s and 100,000 s time points. **C.** The influence of PI(4,5)P_2_ containing liposomes on VP40 PI(4,5)P_2_ binding residues is enlarged. Deuterium incorporation in these regions is plotted in the peptide map to the right of the region of interest. The y-axis is the number of deuterium incorporated into the peptide fragment with respect to time on the x-axis. **D.** The influence of PI(4,5)P_2_ containing liposomes on VP40 PI(4,5)P_2_ residues hypothesized to be important for the stabilization of large oligomers (residues 57-73). Deuterium incorporation in these regions is plotted in the peptide map to the right of the region of interest.

**Figure 9.**
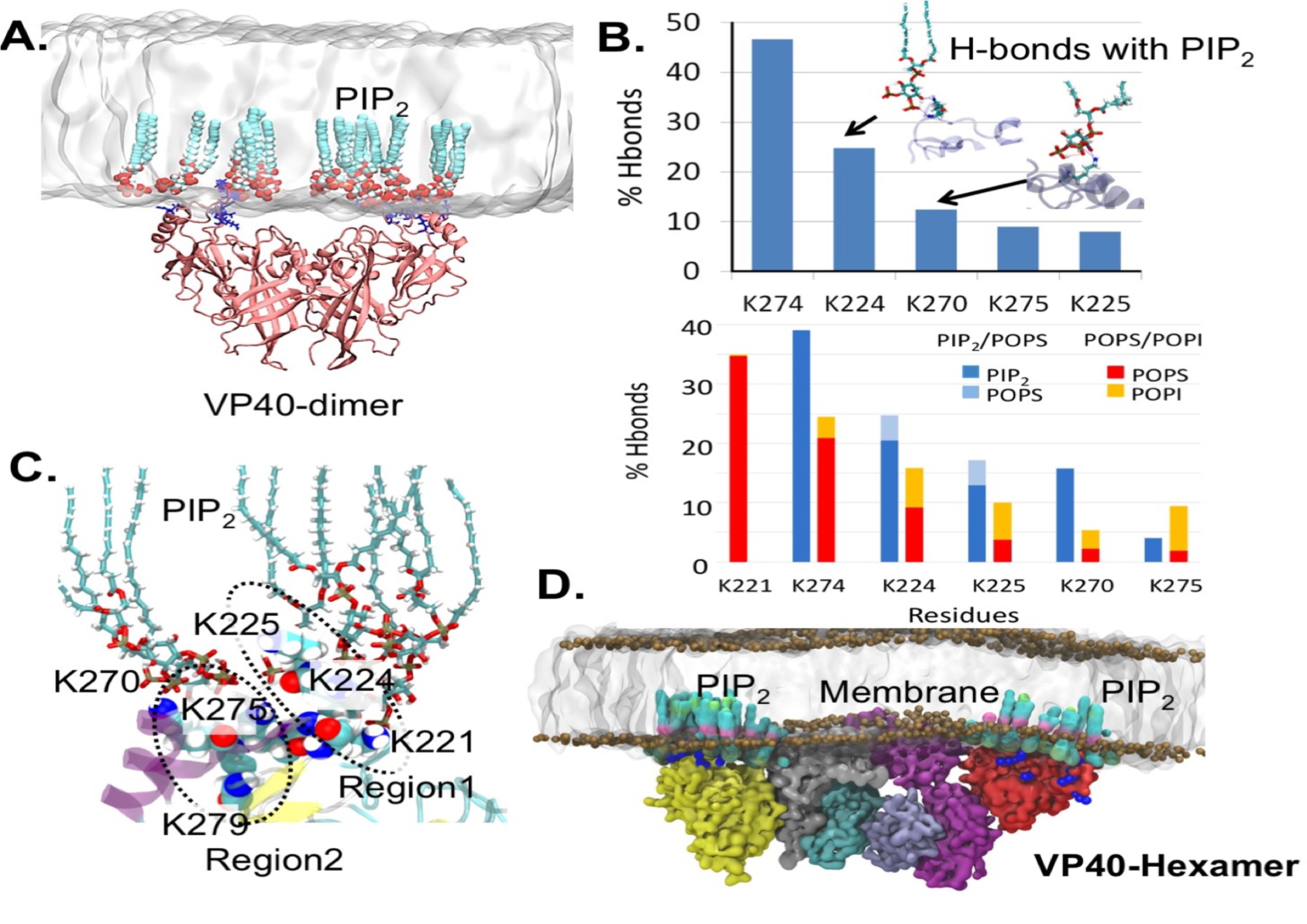
Molecular Dynamics of VP40-PIP_2_ interactions. **A.** VP40 dimer interacting with the PM containing PIP_2_ lipids. **B.** Hydrogen bonding between the CTD-residues and the PIP_2_ head-groups (top panel) and the CTD-residues and PS, PIP_2_ or other lipids (bottom panel) for the VP40 dimer. **C.** Lysine residues in region 1 and region 2 interacting with PIP_2_. **D.** Coarse-grained model showing VP40 hexamer interacting with the PM. The PIP_2_ lipids interacting with the VP40 are highlighted.

Figure 8A shows the percent deuterium incorporation mapped to the hexamer structure while Figure 8B shows the influence of PC, PS, and PI(4,5)P_2_ on the hexamer structure. In the presence of PC, VP40 deuterium exchange is slightly increased compared to without lipid as shown by the light red regions in the NTD. The VP40 structural changes observed through decreased deuterium incorporation after binding PI(4,5)P_2_ were very stable with less than 20% change in deuterium incorporation between the 10 s and 100,000 s time points. The majority of decreases in deuterium incorporation were lost in the PS samples after the 10,000s time point (Figure 8C-D). Again, this data strongly supports a model where PS is important for initial membrane association and VP40 conformational change whereas PI(4,5)P_2_ plays a more predominant role in stabilizing the VP40 oligomers that form.

Deuterium incorporation was reduced at all time points in the regions of the proposed PI(4,5)P_2_ interacting residues. Interestingly, regions containing lys residues in region 220 (Lys^221^, Lys^224,^ Lys^225^) and the region of lysine 270 (Lys^274^, Lys^275^, and Lys^279^) had different deuteration profiles. The 220 lysine residues had a smaller decrease in deuterium incorporation than the 270 lysine residues (Figure 8C). The residues between 221 and 229 have previously been shown to interact with PS, and the HDX supports these observations as the deuterium exchange was reduced for PS containing vesicles slightly greater than that for PI(4,5)P_2_. In contrast, the region containing key Lys residues in PI(4,5)P_2_ binding (Lys^270^, Lys^274^, Lys^275^, and Lys^279^) had a slower rate of exchange with deuterium with either PS or PI(4,5)P_2_ vesicles, with PI(4,5)P_2_ vesicles more significantly reducing deuterium exchange over the time course of the interaction. PI(4,5)P_2_ had a significant affect in slowing the exchange in this CTD region, which correlates with the important role of Lys^274^, Lys^275^, and Lys^279^ in VLP formation and formation of large VP40 oligomers (Figure 6-7).

The most dramatic changes observed were seen in the NTD residues 57-73, a novel site likely important for VP40 oligomer stability. Residues 57-73 are seen packed against the N-N terminal alpha helical interface (Figure 8D). The N-N terminal dimer interface is retained in the hexamer structure; this structure is extremely stable and no deuterium is incorporated even after 100,000 s under all conditions (**Figure S5-S10**). PI(4,5)P_2_ and PS both significantly reduced deuterium incorportation in this N-terminal domain oligomerization interface (residues 57-73), with PI(4,5)P_2_ providing greater stability over the lifetime of the experiments again underscoring the importance of PI(4,5)P_2_ in VP40 hexamers and large oligomers.

#### Molecular Dynamics Section

To gain more molecular insight into PI(4,5)P_2_ coordination by VP40, we performed molecular dynamics simulations of VP40 dimer and hexamers with bilayers containing PI(4,5)P_2_. PI(4,5)P_2_ was shown to cluster at the ends of the VP40 hexamers where interactions between Lys residues in CTD regions 1 and 2 occurred (Figure 9). The H-bonding was monitored over the lifetime of the simulations using an asymmetric bilayer. The lipid composition for the PIP_2_ containing membrane was in the ratio (21:11:32:17:9:10) (CHOL:POPC: POPE: POPS: PIP_2_:PSM) for the inner leaflet of the PM and the outer leaflet had the ratio (19:33:8:3:4:33). Lys^274^ made the most H-bonds over the simulation time with PI(4,5)P_2_ whereas key H-bonds were also found for Lys^224^, Lys^225^, Lys^270^ and Lys^275^. In contrast, Lys^221^ made H-bond contacts with PS only as opposed to PI(4,5)P_2_, which is supported by recent identification of the VP40 PS binding site^22^ suggesting reduction in PI(4,5)P_2_ binding by this mutant may be due to reduced electrostatic interactions. Coarse-grained MD simulations were performed with the VP40 hexamer and an asymmetric bilayer consisting of (POPC: POPE: PSM: POPS: PIP_2_:CHOL) in the ratio (41:8:23:4:4:20) on the upper leaflet and (11:37:5:16:10:21) on the lower leaflet (Fig. 9D). Again, Lys221 made few contacts with PI(4,5)P2 throughout the 1000 ns simulation time whereas significant contacts were observed between PI(4,5)P2 and Lys^274^, Lys^270^, Lys^224^, Lys^225^, and Lys^275^ (**Fig. S14**).

### VP40 budding is reduced when PI(4,5)P_2_ is depleted with small molecule inhibitors

To determine if pharmacological inhibition of PI(4,5)P_2_ synthesis was a feasible approach to inhibit VLP formation, we employed cellular treatments with Phenylarsine oxide (PAO) and Quercetin (Q), which have been used to suppress PI(4,5)P_2_ synthesis^34^. Cell viability was assessed with various concentrations of each drug using the Cell TiterGlo assay kit (see Methods section). Q is a natural product found in many foods^35-39^ and has been reported as a tyrosine phosphatase inhibitor^40^, lipid kinase inhibitor^34^, as well as an anti-microbial, pancreatic lipase inhibitor, and HIV inhibitor^41^. We are interested in the PI5 kinase inhibitory effects of Q, which was shown to inhibit PI(4,5)P_2_ synthesis at the PM^34^. Additionally, HIV-GAG requires PI(4,5)P_2_ for efficient virus particle release and a 40 μM treatment of cells with Q, resulted in 80% inhibition of viral spread^41^. PAO was also shown to inhibit PI(4,5)P_2_ synthesis though inhibition of PI4 kinase IIIα^34^. Previously, we found that a 10 μM, 30 minute PAO treatment significantly reduced VP40 localization from the PM^26^.

Here we tested lower concentrations of these pharamacological agents over a longer treatment time. Confocal imaging of PLCδ-PH with each treatment showed a significant but not complete depletion of PI(4,5)P_2_ from the PM of cells with 500 nM PAO or 100 μM Q compared to the respective DMSO vehicle controls (Figure 10A). The population of cells transfected with VP40 that formed VLPs was also reduced significantly (Figure 10A). Next, VLPs and CL samples were collected and the percent budding was determined with western blot analysis for PAO and Q treatments (Figure 10C and 10D). The VP40 expression was lower in PAO and Q treated cells but the loading control, GAPDH, did not show changes in expression. PAO treated cells had a ~30% decrease in budding compared to the DMSO control, which was not statistically significant over three independent VLP collections. Higher concentrations of PAO are used for more complete PI(4,5)P_2_ depletion in cells, however, these concentrations were toxic to cells over longer treatment times (**Figure S12**). Q treated cells had a statistically significant decrease in budding when compared to the DMSO control. Q treatment in Phase I clinical trials recommended a 1400 mg/m^2^ intravenous treatment^42^. Because PLCδ-PH showed that not all PM PI(4,5)P_2_ is depleted at this concentration of Q, we hypothesize additional Q effects are resulting in the more dramatic decrease in VP40 VLP budding than with PAO treatment. Additional research will prove useful to more closely investigate the mechanism of Q on VP40 function especially as this molecule is a natural product with low toxicity.

**Figure 10:**
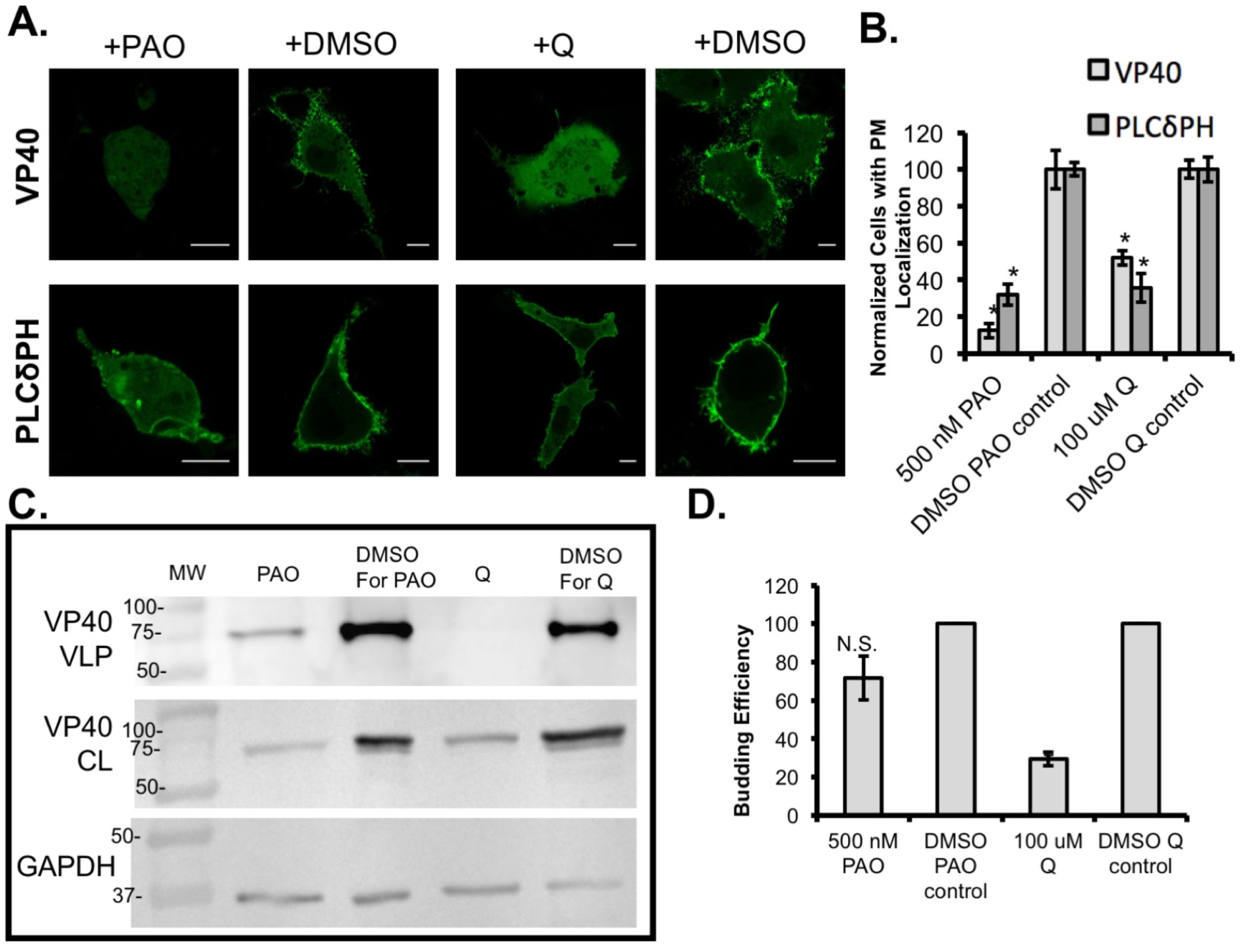
Kinase inhibitors reduce VP40 budding. **A.** Representative images of PLCδPH-EGFP and WT-VP40-EGFP 12 hour post treatment with 500 nM PAO or 100 μM Quercetin (Q). Scale bars are 10 μm. **B.** Cell populations with VLP expression (for VP40) and complete PM localization (for PLCδPH). Values are normalized to cells treated with the equivalent volume of DMSO as a loading control. **C.** Western blots Western blots of WT-VP40-EGFP show VP40 in the VLP and the cell lysate (CL) fractions for each treatment condition, equivalent volumes of DMSO are used as a control for each treatment. GAPDH, was used as a loading control for these experiments. Molecular weight markers are shown for all blots to verify band size for VP40-EGFP and GAPDH. **D.** Budding efficiency is normalized to the DMSO controls. Average values are shown ± the SEM, * P<0.05.

### Discussion

VP40 binds selectively to PI(4,5)P_2_ over the other seven phosphatidylinositol lipids (Figure 2A and 2B). The apparent affinity for PI(4,5)P_2_ containing LUVs is approximately 170 nM (Figure 2C) and this affinity would likely be increased though avidity with VP40 oligomers and likely PI(4,5)P_2_ induced clustering in the cellular environment^28^. Binding of VP40 to PI(4,5)P_2_ is mediated in part by lysine residues in region 1 (K221, K224, K225) and region 2 (K270, K274, K275, K279), which contributed to plasma membrane localization, VP40 oligomerization, and budding (Figures 3-7). Interestingly, these two binding regions have distinct VP40 phenotypes. For example, region 1 lysine residues are required for the formation for oligomers larger than a hexamer whereas region 2 lysine residues harbor more significantly reduced VLP formation but didn’t have as a profound of an effect on the formation of large VP40 oligomers. This data is in agreement with the confocal imaging and assessment of cell populations with VLPs (Figure 4) and SEM imaging (Figure 5).

The HDX MS data confirmed these VP40 binding sites are distinct as they have different deuterium incorporations when bound to PI(4,5)P_2_ liposomes. HDX MS was recently used to study Marburg virus VP40 (mVP40) oligomerization and lipid binding^43^. mVP40 interacts with anionic membranes through electrostatic interactions^44^ and HDX MS confirmed that these interactions occur via CTD residues and that mVP40 undergoes oligomer formation when bound to anionic membranes. In contrast to eVP40, the lifetime of the mVP40 interactions with anionic membranes was relatively short^43^. In region 1 of VP40, PS contributed to a slower exchange rate with deuterium than that of PI(4,5)P_2_ and this region was more important for formation of VP40 oligomers larger than a hexamer. Previously, region 1 was shown to interact with PS^22^ and PS was necessary for formation for formation of VP40 oligomers larger than hexamers^21^. Thus, region 1 may be able to bind PS or PI(4,5)P_2_ with PS providing an interaction that is able to induce VP40 dimer conformational change to a greater extent. More significant exploration of dynamic and structural consequences of lipid head group binding by VP40 will be necessary to delineate the molecular basis of these conformational transitions.

In region 2, PI(4,5)P_2_ slowed exchange more significantly than PS, indicating a more stable VP40 complex. Residues in region 2 had differential effects on VP40 dependent properties depending upon the assay. Lys^274^ seems to be a key PI(4,5)P_2_ interacting residue, which was supported by the MD simulations. Lys275 also made important contributions to PI(4,5)P_2_ binding in *in vitro* and *in silico* assays and like K274A, K275A reduced VP40 oligomerization and VLP formation. The slight differences between K274A and K275A in the assays could be partially attributed to a role of K275A in intermolecular VP40 interactions which decrease the ability of Lys^275^ to be free for membrane binding prior to association with the membrane surface (Fig. S15). K270 and K279 in region 2 also play a significant role in aspects of PI(4,5)P_2_ mediated budding as both K270A and K279A mutants show reduced binding to LUVs and K279A nearly completely abrogated VLP formation. In contrast, K279A large oligomers formed similar to WT suggesting the contacts K279A helps facilitate with the membrane or among VP40 oligomers is critical to budding. Despite a reduction in PI(4,5)P_2_ binding and VP40 oligomerization, K270A was sufficient for approximately the same budding efficiency as WT.

K104A is an interesting mutation as it has VP40 octamer like tendencies in binding (see Figures 3 and **S13**) and forms an abundance of the octamer conformation when purified from bacteria compared to WT VP40 as seen by size exclusion chromatography (data not shown). K104A sits at the N-C terminal domain interface and may be an important residue involved in the unlatching process during self-oligomerization. K104A did have a slight yet non-significant increase in larger oligomers as determined with N&B analysis indicating it may promote larger oligomer formation. Together, our data suggests that K104 is important for stabilization of the dimer N-C terminal domain latch, as there is an increase in binding to neutral lipids with the K104A mutant. Removal of this lysine may result in a destabilized dimer conformation and promote premature oligomerization that results in a significant decrease in budding. Increasing VP40 PM localization and oligomerization does not always indicate that it will promote budding^45^. When VP40 inter-domain dynamics were investigated by GC *et al.*, mutants that promoted PM localization did not show a significant increase in VLP budding^45^. Further investigation of the role of K104 and the potential salt bridge between K104 and D193 may reveal a novel VP40 oligomerization mechanism. We hypothesize that binding to negatively charged PI(4,5)P_2_ or PS at this site could initiate the unlatching of the potential K104-D193 salt bridge.

PAO and Q treatments reduced the overall VP40 expression levels but Q had a significant effect on VLP formation whereas PAO did not. This observation raises the question whether VP40 requires PI(4,5)P_2_ for efficient expression and stability. VP40 enters the nucleus during viral infection^19^ and there is a pool of nuclear PI(4,5)P_2_ that is not completely understood^46^. Alternatively, there is a self-inhibitory effect on VP40 expression if VP40 membrane binding cannot be achieved. The VP40 octamer formation and RNA binding event is critical for viral success, but the VP40 RNA sequence specificity is still unknown. Additionally, the role of VP40 octamer in the nucleus is undetermined but found to be essential for viral success^18^. Additional studies will enhance this mysterious aspect of VP40 function and could provide novel EBOV inhibitory pathways and mechanisms. There are many open questions remaining about the function of the VP40 octamer in the viral lifecycle.

Quercetin treatment shows promise for an Ebola treatment. Could this molecule be adapted for higher potency without an increase in cellular toxicity? Here we found that 100 μM Q treatment and 500 nM PAO treatment result in a decrease in PI(4,5)P_2_ at the plasma membrane in consonance with previous reports^34^. Interestingly, Q had a greater inhibitory effect on budding than PAO. We hypothesize the difference in VLP budding with Q treatment may be from additional inhibitory effects beyond PI(4,5)P_2_ reduction such as tyrosine phosphatase inhibition. Gatto et al. found an 80% reduction in HIV-1 with a 40 μM treatment and found that the inhibitory effects were related to the presence of the C-3 hydroxyl group on Q^41^. Additional research testing other flavonoid molecules against VP40 (BSL 2) or the Ebola virus (BSL 4) could provide powerful insights to viral inhibition. Because Q is a natural product found in several common foods and deemed safe by the FDA as a food additive up to 500 mg per serving, we believe it shows significant promise for testing as a therapeutic against virus replication.

### Methods

#### Molecular Biology

Primers were designed for single and double VP40 mutants and synthesized by Integrated DNA Technologies (Coralville, Iowa). The following primers were used: **K104A**: Forward CCT CTA GGT GTC GCT GAT CAA GCG ACC TAC AGC TTT GAC TC, Reverse GAG TCA AAG CTG TAG GTC GCT TGA TCA GCG ACA CCT AGA GG. **K221A:** Forward GC CCC ATT CTT TTA CCC AAC GCA AGT GGG AAG AAG GGG AAC, Reverse GTT CCC CTT CTT CCC ACT TGC GTT GGG TAA AAG AAT GGG GC. **K224A:** Forward CCC AAC AAA AGT GGG GCG AAG GGG AAC AGT GCC GAT CTA ACA TCT CCG G, Reverse CCG GAG ATG TTA GAT CGG CAC TGT TCC CCT TCG CCC CAC TTT TGT TGG G. **K225A:** Forward CCC AAC AAA AGT GGG AAG GCG GGG AAC AGT GCC GAT CTA ACA TCT CCG G, Reverse CCG GAG ATG TTA GAT CGG CAC TGT TCC CCG CCT TCC CAC TTT TGT TGG G. **K236A:** Forward GCC GAT CTA ACA TCT CCG GAG GCA ATC CAA GCA ATA ATG ACT TCA C, Reverse GTG AAG TCA TTA TTG CTT GGA TTG CCT CCG GAG ATG TTA GAT CGG C. **H269A**: Forward G CCA GAA ACT CTG GTC GCC AAG CTG ACC GGT AAG AAG G, Reverse CCT TCT TAC CGG TCA GCT TGG CGA CCA GAG TTT CTG GC. **K270A**: Forward G CCA GAA ACT CTG GTC CAC GCG CTG ACC GGT AAG AAG G, Reverse CCT TCT TAC CGG TCA GCG CGT GGA CCA GAG TTT CTG GC. **K274A**: Forward G GTC CAC AAG CTG ACC GGT GCG AAG GTG ACT TCT AAA AAT GG, Reverse CCA TTT TTA GAA GTC ACC TTC GCA CCG GTC AGC TTG TGG ACC. **K275A**: Forward G GTC CAC AAG CTG ACC GGT AAG GCG GTG ACT TCT AAA AAT GG, Reverse CCA TTT TTA GAA GTC ACC GCC TTA CCG GTC AGC TTG TGG ACC. **K279A**: Forward CC GGT AAG AAG GTG ACT TCT GCA AAT GGA CAA CCA ATC ATC CC, Reverse GGG ATG ATT GGT TGT CCA TTT GCA GAA GTC ACC TTC TTA CCG G. **K291A**: Forward C CCT GTT CTT TTG CCA GCG TAC ATT GGG TTG GAC CCG, Reverse CGG GTC CAA CCC AAT GTA CGC TGG CAA AAG AAC AGG G. **K224A/K225A:** Forward CCC AAC AAA AGT GGG GCG GCG GGG AAC AGT GCC GAT CTA ACA TCT CCG G, Reverse CCG GAG ATG TTA GAT CGG CAC TGT TCC CCG CCG CCC CAC TTT TGT TGG G. **K274/K275A** (made from K274A template): Forward GGT CCA CAA GCT GAC CGG TGC GGC GGT GAC TTC TAA AAA TGG, Reverse CCA TTT TTA GAA GTC ACC GCC GCA CCG GTC AGC TTG TGG ACC. Site directed mutagenesis was performed using a Quikchange XL cloning kit as described by the manufacturer (Agilent Technologies, Santa Clara, CA). Sanger sequencing was used to verify mutagenesis (Notre Dame Genomics Facility).

#### Protein Purification

VP40-6xHis-pET46 Ek/LIC was expressed in Rosetta BL21 DE3 cells (EMD Millipore Corp, Billerica, MA) as previously described^18^. Briefly, cells were grown to OD_600_ of 0.6 then induced with 1 mM IPTG for 5 hours at RT. Protein was purified with a Ni-NTA affinity column followed by size exclusion chromatography (HiLoad Superdex 200 pg). Protein concentration was determined by BCA assay (Thermo Fisher Scientific, Waltham, MA) and protein was stored at 4°C for up to two weeks.

#### Liposome Pelleting Assay

All lipids were purchased from Avanti Polar Lipids (Alabaster, AL). The liposome pelleting assay was adapted from Julkowska et al.^30^. Lipid films were hydrated with fresh raffinose buffer (250 mM Raffinose pentahydrate, 150 mM NaCl, 10mM Tris, pH 8.0) by thorough heating to 37°C and vortexing. Resuspended lipids were then extruded through a 200 nm filter with an Avanti lipid extruder. Dynamic light scattering (Delsa Nano S particle size analyzer, Brea, CA) was used to confirm liposome size. Liposomes were then diluted and added to the reaction at a final concentration of 1.6 mM lipid and 3.3 uM protein. Reaction mixtures were incubated at room temperature for 30 minutes then centrifuged at 75,000 × *g* for 30 minutes at 22°C.

#### Surface plasmon resonance (SPR)

SPR measurements were made using a Biacore × (GE Healthcare) with an L1 sensor Chip (GE Healthcare). Liposomes were prepared by hydrating lipid films in 10 mM TRIS, pH 7.4 containing 150 mM NaCl followed by extrusion through a 100 nm filter. All experiments were performed at room temperature in 150 mM NaCl, 10 mM TRIS, pH 7.4 using the method previously described in detail^32^. The lipid composition used was POPC:DOPE (80:20) on flow cell 1 and POPC: DOPE: PI(4,5)P2 75:20:5 on flow cell 2. The DPPC:Cholesterol coating had a low response so was not used (data not shown). The apparent affinity (*K*_d_) was determined in Kaleidagraph with the following equation: R_eq_ = R_max_/(1+*K*_d_/[P]) where [P] is protein concentration and R is response units.

#### Cell Culture and transfection

COS-7 cells were maintained in a humidified chamber at 37°C and 5%CO_2_. The cells were kept in DMEM with L-Glutamine, D-Glucose and Sodium Pyruvate (Life Technologies, Carlsbad, CA) with 10% FBS (Sigma, St. Louis, MO) and 1% Penicillin-Streptomycin (Life Technologies) at 37°C, 5%CO_2_. For imaging, cells were seeded in 8-well imaging plates ((MatTek, Ashland, MA) and transfected with Lipofectamine 2000 or Lipofectamine LTX with Plus reagent (Life Technologies), 0.4 ug DNA in optiMEM (Life Technologies) for 12-14 hours. Transfection reactions were scaled up proportionally for scanning electron microscopy and VLP experiments.

#### Confocal Imaging

A Zeiss 710 laser scanning microscope was used to image EGFP-VP40 phenotype in COS-7 cells. A 488 nm laser was used to excite EGFP. All EGFP-VP40 expressing cells were counted as either presenting VLPs or not were counted. VP40-EGFP mutants were compared to WT. Each construct was counted and imaged over at least three independent experiments. For number and brightness experiments, an Olympus FV1000 instrument was used according to the procedure explained in depth previously^47^. Data analysis was performed in SimFCS software (Globals Software, Irvine, CA) as previously described in detail^47^.

#### Scanning Electron Microscopy

Transfected cells were scraped from plates 12-14 hours post transection as described previously in detail^31^. Images were acquired on a Field Emission Scanning Electron Microscope Magellan 400 (FEI, Hillsboro, OR) at the Notre Dame Integrated Imaging Facility (NDIIF).

#### VLP Collection and Western Blot Analysis

VLPs were harvested 24 hours post transfection as described previously in detail^31^. Briefly, VLPs were washed from cells with 1X PBS, and applied to a sucrose cushion. Samples were centrifuged at 100,000 × *g*, at 4°C for 2 hours. During the centrifugation, cells were scraped from plates and lysed with RIPA buffer. Soluble protein was isolated from cell lysate via centrifugation at 17,000 × *g* at 4°C for 15 minutes, followed by removal of the soluble fraction of samples. Samples were then stored at −80°C. A BCA assay was used to determine protein concentration of all samples, 15 ug of each cell lysate was loaded for SDS PAGE and Western Blot. A proportional volume of the VLP sample was loaded for SDS PAGE and Western Blot. For VP40 detection, mouse anti-EGFP (F56-6A1.2.3, ThermoFisher Scientific) primary antibody and sheep anti-mouse HRP (AB 6808, Abcam, Cambridge, United Kingdom) secondary antibody was used. For GAPDH detection, an anti-GAPDH antibody was used (AB 8245, Abcam) and sheep anti mouse HRP (AB 6808, Abcam) secondary antibody was used. ECL blotting substrate was used according to the manufactures instructions (Thermo Scientific). Blots were imaged on a GeneGnome (Syngene, India). ImageJ was used to quantify band density and determine budding efficiency.

#### Hydrogen Deuterium Exchange Mass Spectrometry (HDX MS)

HDX MS experiments were performed as described in detail previously with Marburg VP40 43. For Ebola VP40, multilamellar vesicles (MLVs) of the following compositions were used: PC (50% POPC, 50% DOPE), 12.5% PI(4,5)P_2_ (47.5% POPC, 40% DOPE, 12.5% PI(4,5)P_2_), and 60% POPS (30% POPC, 10% DOPE, 60% POPS). 3 ug protein and 2.8 mM MLV were incubated 30 minutes prior to the HDX experiments. HDX data was analyzed as previously described in detail 43 and mapped to eVP40 dimer (4ldb) and hexamer (4Ldd with modeled C-terminal domains courtesy of Dr. Prem Chapagain^28^).

#### Cell Viability Assay

Cell viability was determined using the CellTiter Glo assay (Promega, Madison, WI) according to the manufactures instructions. Cells were treated with indicated therapeutics for 24 hours in optiMEM (Life Technologies). The viability assay was performed at least two independent times for each drug concentration or equivalent volume of vehicle, DMSO. Drug concentrations that allowed for 80% cell viability or more were tested further in imaging and VLP collection experiments.

#### Pharmacological Treatments

Phenyl arsine oxide (PAO) (Sigma) and Quercetin (Q) (Sigma) stocks were prepared in DMSO and diluted in OptiMEM (Life Technologies) prior to use. For imaging experiments, cells were treated with PAO or Q for 14 hours after being added 4 hours post transfection. For VLP collection experiments, cells were transfected with Lipofectamine LTX (Life Technologies) then treated with PAO or Q for 20. In both experiments, transfection media was removed with aspiration and replaced with drug or DMSO containing optiMEM. The equivalent volume of DMSO was added as a vehicle control for PAO and Quercetin imaging and VLP collection experiments. Each imaging experiment and VLP collection was performed three independent times.

#### Reproducibility and Statistics

At least three experimental replicates were performed for all experiments unless otherwise noted. A two tailed students T-test was used to determine statistical differences. A P value of 0.05 or less was considered statistically significant. Error bars are shown as the standard error of the mean in all graphs unless otherwise noted.

#### All-atom simulations of VP40 Dimer and Plasma Membrane

The crystal structure of VP40 dimer was obtained from the Protein Data Bank (PDB ID: 4LDB). The protein and plasma membrane (PM) complex was set up^48^ using the Charmm-Gui web server^49^. The PM systems consisted of different types of lipids: 1-palmitoyl-2-oleoyl-sn-phosphatidylcholine (POPC), 1-palmitoyl-2-oleoyl-sn-phosphatidyl-ethanolamine (POPE), 1-palmitoyl-2-oleoyl-sn-phosphatidyl-serine (POPS), palmitoylsphingomyelin (PSM), 1-palmitoyl-2-oleoyl-sn-glycero-3-phosphoinositol (POPI), palmitoyl-oleoyl-phosphatidylinositol-(4,5)-bisphosphate (PIP_2_) and cholesterol (CHOL). Two different systems were constructed: one with PIP2 and another with POPI. The lipid composition for the PIP_2_ containing membrane was in the ratio (21:11:32:17:9:10) (CHOL:POPC: POPE: POPS: PIP2:PSM) for the inner leaflet of the PM and the outer leaflet had the ratio (19:33:8:3:4:33). There were 442 lipids in the outer leaflet and 451 on the inner leaflet. A similar membrane was constructed containing POPI with a lipid composition ratio of (21:11:32:17:9:10) (CHOL: POPC: POPE: POPS: POPI: PSM) for the inner leaflet and (19:32:8:3:4:34) for the outer leaflet. There were 149 lipids in outer leaflet and 151 in the inner leaflet for this system.

Both dimer-membrane systems were solvated using TIP3 water molecules. The total system consists of around 361000 atoms (PIP_2_ setup) and around 121,000 atoms for (POPI set up). We performed all-atom molecular dynamics simulations using NAMD2.10^50^ with the CHARMM36 force field for both systems. The particle mesh Ewald (PME) method was used in order to treat long range electrostatic interaction and the SHAKE algorithm was employed to constrain hydrogen-containing covalent bonds. The system was minimized for 10,000 steps followed by a six step equilibration process. Pressure was controlled using a Nose-Hoover Langevin-piston method. Langevin temperature coupling with a coefficient friction coefficient of 1 ps^−1^ was used to control the temperature and a 2 fs time step was used for production runs.

#### Coarse-grained VP40 Hexamer and Plasma Membrane

The crystal structure of the VP40 hexamer was obtained from the Protein Data Bank (PDB: 4LDD). Missing residues were added using the Modeller software package. The plasma membrane and hexamer system was built^28^ using the Charmm-gui webserver^49^. The composition of the various lipids (POPC: POPE: PSM: POPS: PIP_2_:CHOL) were in the ratio (41:8:23:4:4:20) on the upper leaflet and (11:37:5:16:10:21) on the lower leaflet. Since the total number of atoms was greater than 600,000 atoms, a coarse grained (CG) model was constructed to reduce the number of particles so that computational simulations of hundreds of nanoseconds were feasible. For CG simulations, we used the Martini 2.0 force field, where four heavy (non-hydrogen) atoms were mapped onto a single bead, excluding residues with aromatic side chains. The lipids were modelled as in Gc et al.^28^ and Van Der Spoel et al.^51^. The system was solvated using standard Martini water beads and was neutralized in 0.15 M NaCl. The total number of particles for the CG system was 52827.

The coarse-grained molecular dynamics simulation was performed using Gromacs 5.1.1^51,52^. The simulation was performed with 15 fs time steps. The Lennard-Jones and Coulomb potential had cutoffs at 11 Å. In order to keep track of particles, a Verlet neighbor scheme was used and Coulomb interactions were treated using a reaction field. The LINCS algorithm was used to control stiff bonds. The pressure was maintained at 1 bar using a Berendsen barostat for equilibration runs and a Parrinello-Rahman barostat during production runs. Pressure coupling between the protein and membrane was semi-isotropic with a compressibility of 3×10^−4^ bar^−1^ and the temperature was maintained at 303 K. Temperature coupling was done using a velocity rescale algorithm.

## Acknowledgments

This research was supported by NIH AI081077 (R.V.S) and1U19AI117905, R01GM020501, R01NS070899, and R01GM121964 (S.L.). K.A.J would like to thank the NDIIF and Dr. Sergi Rouvimov and Dr. Tatyana Orlova for helpful discussions in TEM and assistance in SEM, respectively. The confocal and multiphoton imaging studies were supported by the Indiana University School of Medicine-South Bend Imaging and Flow Cytometry Core.

